# Targeting PrfA from Listeria monocytogenes: a computational drug repurposing approach

**DOI:** 10.1101/2023.12.10.570966

**Authors:** Xabier Arias Moreno

## Abstract

Humans can develop listeriosis by ingestion of foods contaminated with *Listeria monocytogenes,* an opportunistic gram-positive ubiquitous bacterium. Whilst the non-invasive form of listeriosis may be asymptomatic or cause mild flu-like symptoms, the invasive form of listeriosis is life-threatening and is associated with high hospitalization and fatality rates. Current antibiotic-based therapies are still effective against listeriosis. However, multi-drug resistant *Listeria monocytogenes* strains have already been identified, which represents a new risk in the treatment of invasive listeriosis. Therefore, it is increasingly urgent to identify new compounds that do not target the conventional biochemical pathways disrupted by current antibiotics. Positive Regulatory Factor A (PrfA) is a well-studied transcriptional factor in *Listeria monocytogenes* that is responsible for activating a plethora of virulence factors. Targeting virulence factors is a promising strategy that is being considered to combat bacterial infections, hence targeting PrfA is both logical and attractive. In the present computational drug repurposing approach, a complete FDA-approved drugs dataset of more than 700 compounds was virtually screened by docking the drugs against the structure of PrfA. Three of the most promising top-scored FDA drug candidates were then simulated complexed to PrfA. Data from Molecular docking and Molecular Dynamics simulations suggest that Dutasteride and Solifenacin may bind PrfA, and as a result, might exhibit inhibitory activity against *Listeria monocytogenes*. The use of Dutasteride and Solifenacin is safe in humans since they have been used in the treatment of benign prostatic hyperplasia and androgenic alopecia, and urinary incontinence respectively for decades. Their unique chemical scaffolds may represent valuable starting points for the rapid development of disruptive novel listeria-specific drugs that will be soon needed to combat multi-drug resistant *Listeria monocytogenes* strains.

## 1. INTRODUCTION

*Listeria monocytogenes* (*L. monocytogenes*) is a facultative anaerobic, ubiquitous gram-positive bacterium with a remarkable ability to grow at low temperatures and to resist low water activity, low pH and high salt environments [1,2]. *L. monocytogenes* is also capable of forming biofilms, which further helps the microorganism withstand the most harsh environments [3]. Ingestion of foods contaminated with *L. monocytogenes* usually produces mild flu-like symptoms such as nausea, vomiting and diarrhea. In contrast, the invasive form of listeriosis can develop into sepsis, meningitis and encephalitis, which are all life-threatening conditions [4]. The elderly, immunocompromised individuals, pregnant women, newborns, and fetuses are particularly at risk of developing the invasive form of the disease [5]. Foods commonly involved in listeriosis cases are raw or processed vegetables, unpasteurized milk and cheese, ice creams, raw or processed fruits, raw or smoked fish and seafood, and Ready-To-Eat (RTE) processed foods [6]. Even though the prevalence of listeriosis is very low, the fatality rate of the invasive form is 20 to 30% [5], making *L. monocytogenes* one of the most dangerous foodborne pathogens. In 2021, the number of confirmed cases of human listeriosis in the European Union was 2,183 and the associated fatality rate, among the cases with known outcomes, was 13.7% [7]. Listeriosis outbreaks, which are not infrequent, can sometimes affect a large number of individuals. In one of the most deadliest and recent outbreaks, more than 200 people died in South Africa [8]. Due to the long incubation period for both invasive and pregnancy-associated listeriosis [9], identification of the contaminated food source during an outbreak is extremely challenging.

The current gold standard antibiotic therapy in the treatment of invasive listeriosis is based on the administration of penicillins such as aminopenicillin or benzylpenicillin, alone or in combination with aminoglycosides, usually gentamicin [10]. Treatment of *L. monocytogenes* with these antibiotics is still effective. However, multi-resistant *L. monocytogenes* strains have been isolated and identified in human and food samples [11–16]. These multi-resistant strains pose a high risk in the treatment of invasive listeriosis [17]. Therefore, the development of new drugs targeting unconventional biochemical pathways and bypassing antibiotic resistance is of utmost importance [17]. Traditional drug development is a time-consuming and expensive process as putting a new drug on the market is estimated to cost more than one billion dollars with an average of more than 15 years from hit identification to drug commercialization [18]. Of 5,000 compounds that enter the preclinical phase, only five reach the clinical trials phase, and only one receives approval for therapeutic use [19]. In contrast, drug repurposing, also known as drug repositioning, is a cost-effective drug development alternative [9]. Screening drugs that have already been approved for other indications represents an attractive strategy for speeding up the identification and development of new applications for drugs already on the market. Countless examples of successful drug repurposing applications exist, but perhaps some of the most recent and significant are related to COVID-19 [20]. Nirmatrelvir, Remdesivir, and Molnupiravir, which all are currently recommended by WHO in the treatment of non-severe COVID-19 at highest risk of hospitalization, are repurposed drugs [21,22]. Nirmatrelvir, which was initially developed to inhibit the main protease of SARS-COV-1 virus, was recently found to also inhibit the main SARS-COV-2 protease, M^pro^ [23]. Remdesivir, which was developed to treat hepatitis C, was recently found to inhibit the viral RNA-dependent RNA polymerase (RdRp) of SARS-COV-2 [24]. Molnupiravir, a nucleoside derivative that was originally developed to treat the Flu, was found to inhibit the replication of SARS-COV-2 [25].

Positive Regulatory Factor A (PrfA) is a transcriptional factor in *L. monocytogenes* that is responsible for the transcription and activation of multiple key virulence factors [26]. PrfA is considered the main regulator of *L. monocytogenes’* virulence and its role in bacterial propagation inside the host is well-documented [27,28]. At the molecular level, PrfA is a homodimer where each of its 27 kDa monomers (chain A and chain B), is composed of 237 amino acids (**Figure 1A**) [29]. High-affinity DNA binding of PrfA only occurs after allosteric binding of reduced Glutathione (GST) to each of its chains [30]. GST binds to a huge hydrophobic pocket called the coactivator binding site (**Figure 1B**), which is also known as the tunnel site. The coactivator binding sites in chain A and chain B are sometimes referred as site A_I_ and Site B_I_ respectively [29], and is is the preferred terminology used in the present manuscript. The N-terminal domain of PrfA is a β barrel and is linked to the C-terminal domain by a long α-helix. The C-terminal domain is an α**/**β domain and bears the Helix-Loop-Helix (HLH) motif that directly interacts with the PrfA boxes scattered across *L. monocytogenes*’ genome [29]. In recent years, several synthetic small molecule PrfA inhibitors have been identified and characterized in detail, such as C01, C16 (**Figure 1C**), IWP-2 and PS900 among others [31–34]. These molecules, which bear a ring-fused 2-pyridone scaffold, are considered competitive inhibitors of GST. Although this strategy of targeting the virulence of the pathogen is promising, it is non-conventional, as the traditional biochemical pathways disrupted by antibiotics are not targeted [33,35–37]. Additional studies will certainly be needed to confirm the inhibitory and bacteriostatic activity of these novel PrfA inhibitors in pre-clinical studies [31]. In any case, the benefits of this strategy are extraordinary as completely novel biological entities can be identified [38]. These PrfA inhibitors could be used as starting points in the development of novel drugs that, in principle, will be effective against multi-drug resistant *L. monocytogenes* strains. An additional benefit of using these inhibitors, and contrary to what is observed with antibiotics, is that they will help preserve the microbiome of the host.

**Figure 1.**
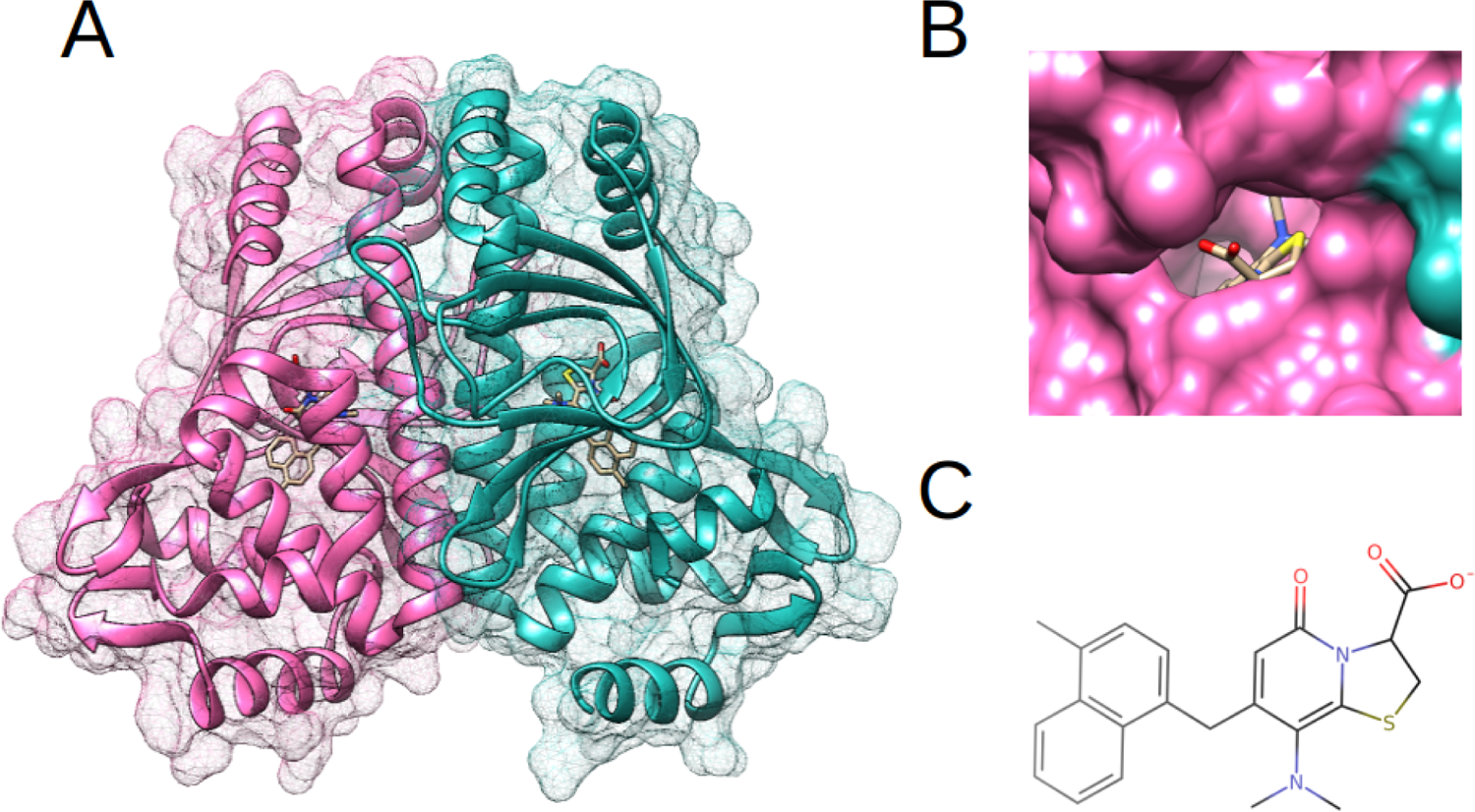
PrfA transcriptional factor and C16 inhibitor. A) C16:PrfA complex as seen in the 6EXL PDB file. Two C16 molecules are seen, one inside chain A and one inside Chain B of PrfA. PrfA chains are visible as ribbons with a mesh surface transparency whilst C16 inhibitors are visible as sticks. PrfA chains A and B are depicted in light pink and light sea green respectively. B) Coactivator binding site in PrfA. The C16 inhibitor is seen inside site A_I_ of PrfA. PrfA is visible as solid molecular surface whilst C16 inhibitor is visible as sticks. PrfA chains A and B are depicted in light pink and light sea green respectively. Figures 1A and 1B were prepared and rendered on Chimera. C) Molecular structure of C16 inhibitor.

In a previous computational work, we described several potential small molecule PrfA inhibitors, such as P875 and P584, that were identified after applying a combined ligand-based virtual screening, structure-based virtual screening, and Molecular Dynamics (MD) strategy [39]. In contrast, the goal of the present computational drug repurposing approach was to identify FDA drugs with potential PrfA inhibitory activity. To achieve this, a strategy combining structure-based virtual screening and MD simulations has been utilized (**Figure 2**). First, an FDA-approved drugs dataset with more than 700 compounds was docked against site A_I_ of PrfA using PyRx. Then, the 25 top-scored drugs were docked against both site A_I_ and site B_I_ of PrfA using Autodock Vina and a combined average binding energy was calculated for each compound. Finally, the three FDA drugs with the highest average binding energies were simulated complexed to PrfA using Gromacs. The molecular docking and the all-atom 100-ns MD simulations suggest that Dutasteride and Solifenacin drugs may bind PrfA and, as a result, might exhibit inhibitory activity against *L. monocytogenes*.

**Figure 2.**
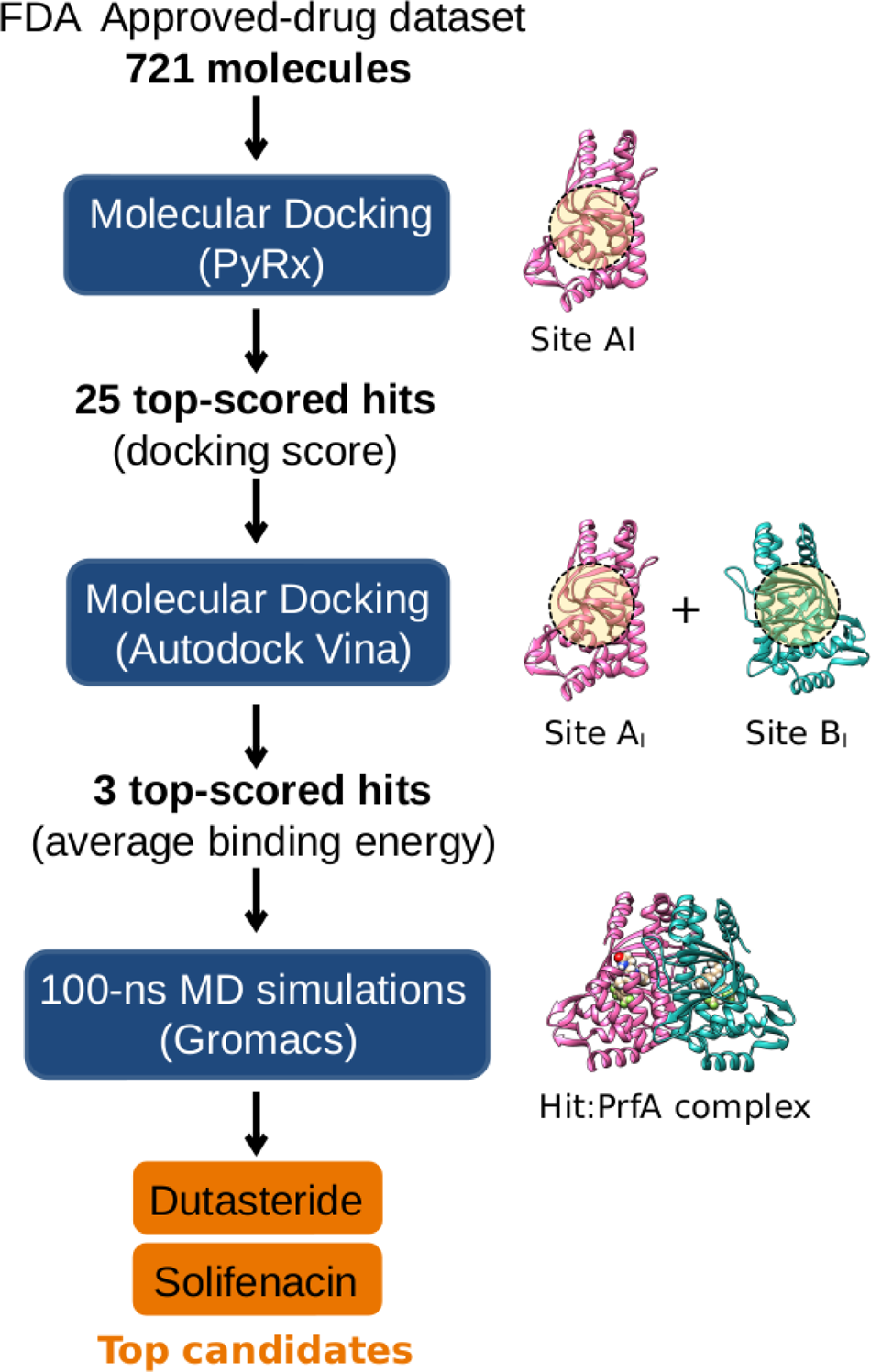
Schematic representation of the work-flow algorithm used in the present computational repurposing work. A total of 721 molecules were docked on Site A_I_ of PrfA using PyRx. The 25 top-scored compounds were further docked on both Site A_I_ and Site B_I_ of PrfA using Autodock Vina. The three top-scored compounds were complexed to PrfA and their MD simulations were performed on Gromacs. The top candidates of this strategy are Dutasteride and Solifenacin. Figures were prepared and rendered on Chimera. For more information see text.

## 2. MATERIALS AND METHODS

### 2.1 Computational Drug Repurposing strategy

The overall drug repurposing strategy utilized in the present work is summarized in **Figure 2**. Briefly, a total of 721 molecules derived from a FDA-approved drugs dataset were first docked against site A_I_ of PrfA using PyRx 0.8. Then, the compounds were sorted according to their docking scores. Next, the 25 top-scored hit compounds were docked against site A_I_ and site B_I_ of PrfA using Autodock Vina 1.1.2. Then, the 25 hit compounds were sorted according to their calculated average binding energies. Lastly, 100-ns MD simulations of the three top-scored hit compounds complexed to PrfA were performed using Gromacs 2021. A detailed explanation of each of the steps is described in the following sections.

### 2.2 Selection of the PrfA structure and the Positive control

For this work, C16 compound inhibitor (**Figure 1C**) was treated as the positive control molecule instead of more recently described PrfA inhibitors such as IPW-2 and PS9000. C16 compound had already been used as a positive control in our previous computational work, where it was shown to produce a very stable PrfA complex, and therefore was decided to give it the same consideration [39]. Therefore, the PrfA structure utilized in the present work was extracted from the 6EXL PDB file (**Figure 1A**), which is the same exact structure utilized in our previous work. In this structure, PrfA appears bound to C16 compound. 6EXL structure was chosen over 6EXK, the other available C16:PrfA complex structure, because in 6EXL one of the HLH motifs is well-resolved while in 6EXK both HLH motifs are missing from the PrfA structure. 6EXL PDB file was downloaded from the RCSB PDB Protein Data Bank [40] and its structure was prepared as described before [39]. For the present work, water molecules were also deleted. The corresponding PDBQT file was generated using Autodock Tools 4.2 [41]. Molecular docking was performed using PyRx 0.8 [42] and Autodock Vina 1.1.2 [43]. The compounds were docked against site A_I_ of PrfA (**Figure 1B**) [18] on PyRx and against both site A_I_ and site B_I_ of PrfA on Autodock Vina, as done previously [39].

### 2.3 Structure-based Virtual Screening using PyRx

The FDA-approved drugs dataset was downloaded from the ZINC15 database [44] as a MOL2 file. Virtual screening was performed using PyRx 0.8 [42], an open-source software that uses the Autodock Vina algorithm [42]. First, compounds from the drugs dataset were brought to the lowest energy level by performing 200 total minimization steps under the Universal Force Field. Then, each of the compounds was extracted and transformed automatically into individual PDBQT files using Open Babel implemented on PyRx. Virtual Screening was performed in a grid box of 18.0 Å x 25.0 Å x 20.0 Å that was centered in X=-24.6, Y=-17.08 and Z=20.4 coordination space, which corresponded to site A_I_ in PrfA. The Vina protocol was run at an exhaustiveness of 16.

Validation of the virtual screening using PyRx was done by including C16 compound within the FDA-approved drugs dataset. First, the coordinates of the two top-scored C16 molecule poses obtained on PyRx were transformed from PDBQT to PDB files. Then, the atom numbering of the two C16 molecules within the two PDB files was unified on a text editor. Finally, the Root Mean Standard Deviations (RMSD) between the crystallographic and the docked poses obtained on PyRx were calculated using VMD 1.9.3 [45].

### 2.4 Molecular docking using Autodock Vina

The 25 top-scored compounds obtained on PyRx were downloaded from the PubChem database [46] as 3D SDF files together with their main impurities, metabolites and isomers available for each of the compounds. The protonation state of all the downloaded compounds was set at pH 7.4 on OpenBabel 3.1.1 [47]. All compounds and their impurities, metabolites and isomers were carefully prepared for molecular docking using Autodock Tools 4.2, where the PDBQT files were obtained, as done before [39]. The hydrogen content, atom coordinates, molecule charge and bond torsions were carefully checked on UCSF Chimera 1.17.1 [48] and on a text editor. Then, all molecules were docked against both site A_I_ and site B_I_ of PrfA using Autodock Vina 1.1.2 [43]. Virtual Screening was performed in a grid box of 35.0 Å x 35.5 Å x 35.0 Å that was centered in X=-24.4, Y=-13.5 and Z=31.1 (site A_I_) and in X=-0.4, Y=-9.3 and Z=29.0 (site B_I_) coordination spaces. The Vina protocol was run at an exhaustiveness of 16. A minimum of three individual molecular docking were performed for each of the molecules. The highest docking score result was considered. For each compound, an average binding energy was calculated. This average binding energy corresponds to the average Vina docking score energies obtained after docking against each of PrfA’s chains. Again, the C16 compound was included in the dataset for validation purposes. The molecular structures of the 10 top-scored compounds and the C16 compound were generated using Marvin JavaScript 23.11.0 [49].

### 2.5 Selection of the PrfA structure and hit molecule poses for MD simulations

The three top-scored hit compounds obtained after performing molecular docking on Autodoc Vina were simulated complexed to PrfA. PrfA:C16 complex was also simulated and was treated as a positive control. The PrfA structure that was used in the MD simulations was the same 6EXL structure that was used during the molecular docking. Selection of the adequate compound poses obtained in the molecular docking for the MD simulations was trickier. In this respect, it would have been tempting to have just selected the top-scored pose of the hit compounds, i.e. pose 1, obtained on each PrfA’s chains. However, pose 1 conformation from chain A and chain B did not always match.

As a general rule, both pose 1 conformations obtained from chain A and chain B were only selected if they happened to have an identical or very similar conformation and orientation inside the PrfA structure. In the rest of the cases, the conformation of pose 1 with the higher docking score prevailed. Then, the homologous conformation pose obtained on the other PrfA chain was identified and selected. As a consequence, some of the complexes were simulated with one of the compound hit molecules that were not pose 1. In a homodimeric symmetric protein system like PrfA, simulation of a PrfA complex with a compound molecule in chain A and its flipped version in chain B would certainly have been unrealistic.

### 2.6 MD simulations using Gromacs

MD simulations of PrfA complexed to the three top-scored compounds were carried out in Gromacs 2021 [50] on an in-house Linux-based desktop computer. Validation of the MD simulations was done by simulating the PrfA:C16 complex, as done previously [39]. The topology and coordinate files of the ligands were generated on the SwissParam web server [51]. The total charge of the molecules on the ITP files was individually checked on a text editor. All the simulations were carried out on the CHARMM 36 all-atom force field. A 10 Å dodecahedron box was filled with CHARMM-modified TIP3 water model molecules. The cut-off for short-range electrostatic and van der Waals interactions was set to 14 Å. PME method was used for long-range electrostatic interactions. Cl^-^ anions were added to equilibrate the system. Energy minimization was performed using the steepest descent algorithm with an energy convergence cut-off of 1,000 kJ/mol. Periodic Boundary Conditions were applied in all directions. Temperature equilibration was performed for 0.5 ns, where position restraints were applied for both protein and ligands. Constraints were applied for all hydrogen atoms. The time step was set to 1 fs and the temperature coupling was maintained with V-rescale. Pressure equilibration was performed for 0.5 ns, where position restraints were applied for both protein and ligands. Constraints were applied for all hydrogen atoms. Time step was set to 1 fs and the pressure coupling was maintained with C-rescale. Productive simulations were performed for 100-ns with a time step of 1 fs at a constant 1 atm pressure and 310 K with the same thermostat and barostat used during the equilibration step. Constraints were applied for all hydrogen atoms. Three independent MD simulations were performed for each of the PrfA complexes with newly assigned initial velocities. Backbone RMSD of PrfA chains and RMSD of the compound molecules were calculated using a built-in utility installed on Gromacs. RMSD graphs were prepared on Grace 5.1.25. The average and Standard Deviation (SD) RMSD values for each of the simulations were calculated on a spreadsheet editor.

## 3. RESULTS

### 3.1 Structure-based Virtual Screening using PyRx

Of the 1,345 drug molecules within the FDA-approved drugs dataset, only 721 molecules were successfully converted into PDBQT files. These 721 molecules, which represents the 54% of the molecule library size, were docked against site A_I_ of PrfA using PyRx. The 25 top-scored hit compounds spanned docking scores from − 10.8 kcal/mol to −12.4 kcal/mol (**Table 1**). The C16 compound produced a docking score of −11.1 kcal/mol. Interestingly, more than half of the 25 compounds have either a steroid scaffold or are derived from the cholesterol molecule. On the one hand, the molecules with steroid scaffold are Dutasteride, Progesterone, Exemestane, Prasterone, Clocortolone, Medroxyprogesterone acetate, Fluometholone, Oxymetholone, Oxandrolone, Estrone sulfate, and Desonide. On the other hand, the three molecules derived from the cholesterol molecule are Ergocalciferol, also known as Vitamin D_2_, and Vitamin D analogs Calcipotriol and Paricalcitol.

**Table 1.** 25 top-scored compounds obtained after performing Virtual Screening on PyRx. FDA-approved drug molecules were docked against Site A_I_ of PrfA. The ZINC ID, drug name and PyRx docking score energy is shown for each of the compounds. The 26 top-scored molecules are shown due a docking scoring tie between the 25^th^ and 26^th^ compound.

| **Top-score Rank** | **ZINC ID** | **Drug name** | **PYRX Score** |
| --- | --- | --- | --- |
| **1** | 1164737 | Atovaquone | -12.1 |
| **2** | 3480647 | Enzalutamide | -12.0 |
| **3** | 3932831 | Dutasteride | -11.7 |
| **4** | 403566 | Praziquantel | -11.7 |
| **5** | 4428529 | Progesterone | -11.6 |
| **6** | 1539579 | Bexarotene | -11.6 |
| **7** | 3973334 | Exemestane | -11.5 |
| **8** | 3936683 | Solifenacin | -11.4 |
| **9** | 3807917 | Prasterone (DHEA) | -11.3 |
| **10** | 3816514 | Rolapitant | -11.3 |
| **11** | 2198145 | Clocortolone | -11.3 |
| **12** | 5029557 | Medroxyprogesterone | -11.2 |
| **13** | 6094354 | Palonosetron | -11.2 |
| **14** | 1189125 | Fluorometholone | -11.2 |
| **15** | 4629876 | Ergocalciferol | -11.1 |
| **16** | 968264 | Cyproheptadine | -11.0 |
| **17** | 1250306 | Trospium | -11.0 |
| **18** | 1189124 | Oxymetholone | -11.0 |
| **19** | 3802417 | Alvimopan | -10.9 |
| **20** | 3813047 | Oxandrolone | -10.9 |
| **21** | 3876186 | Estrone sulfate | -10.9 |
| **22** | 4212851 | Desonide | -10.8 |
| **23** | 3921872 | Calcipotriol | -10.8 |
| **24** | 1000709 | Glimepiride | -10.8 |
| **25** | 75126 | Ondansetron | -10.8 |
| **26** | 1391194 | Paricalcitol | -10.8 |
| **N/A** | **C16** | **Control** | **-11.1** |

### 3.2 Validation of the Virtual Screening using PyRx

Validation of the virtual screening using PyRx was performed by including the C16 compound within the FDA-approved drugs dataset. The obtained docking score was −11.1 kcal/mol for C16 pose 1 and −11.0 kcal/mol for C16 pose 2. The RMSD between the two poses was 2.38 Å. When both poses were compared to the crystallographic C16 conformation seen in the 6EXL crystal structure on chain A of PrfA, the obtained RMSD values were 0.24 Å and 2.35 Å for pose 1 and pose 2 respectively (**Figure 3**). The obtained RMSD value of 0.24 Å is a very good indicator that adequate docking parameters were utilized on PyRx.

**Figure 3.**
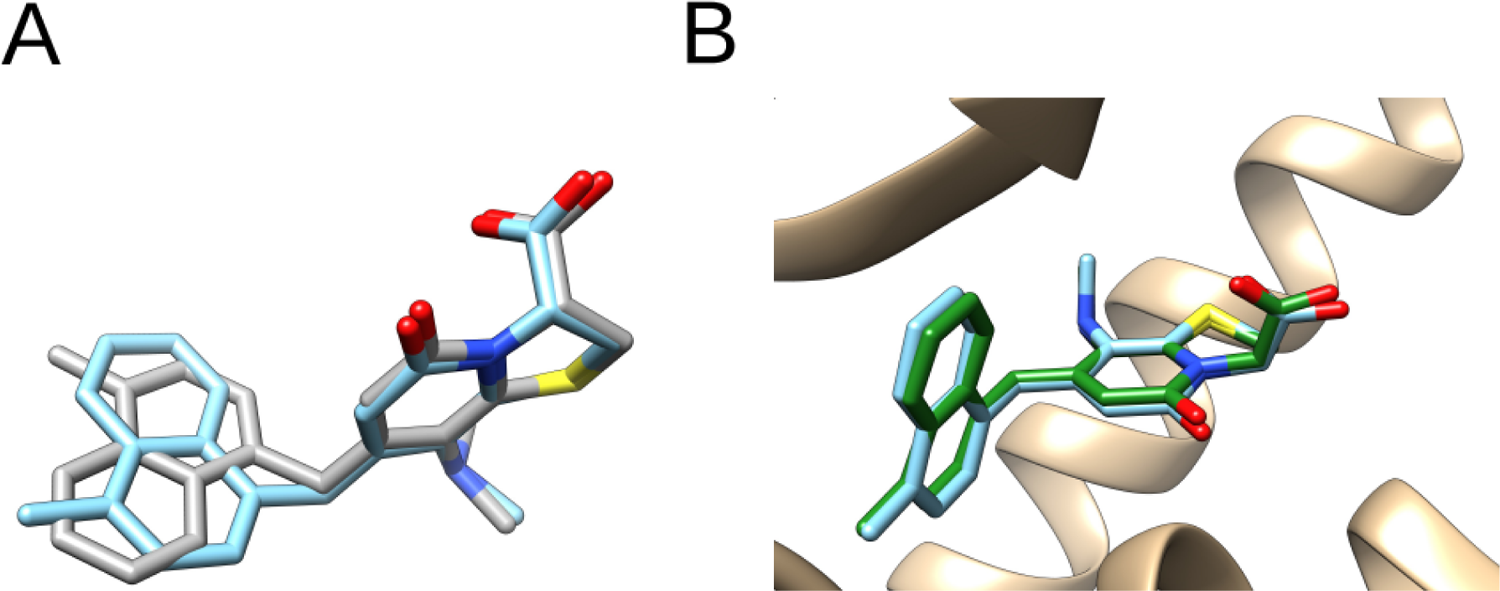
Validation of virtual screening on PyRx (site A_I_). A) Superposition of pose 1 and pose 2 of C16 compound obtained on PyRx. B) Superposition of C16 pose 1 and its corresponding crystallographic conformation seen in the 6EXL pdb. C16 molecules are represented in sticks with colored heteroatoms. C16 pose 1 is depicted in light blue, C16 pose 2 in light gray and the crystallographic C16 pose in green. PrfA is represented in ribbon and depicted in light brown. Figures were prepared and rendered on Chimera.

### 3.3 Molecular docking using Autodock Vina

The 25 top-scored compounds obtained using PyRx were further evaluated by docking them and their impurities, metabolites, and isoforms against site A_I_ and site B_I_ of PrfA using Autodock Vina. For each compound, an average binding energy was calculated as explained before [39] and in section 2.3. The 10 top-scored compounds were sorted according to the obtained average binding energies (**Table 2**). Of note, only the highest average binding energy of each compound or its related impurities, metabolites and isomers is shown. The obtained average binding energies spanned energies from −10.7 kcal/mol to −12.2 kcal/mol, which were considerably higher than the average binding energy obtained for the C16 compound, which was −10.4 kcal/mol. Interestingly, IWP-2 inhibitor [33] produced the same average binding energy as C16 (not shown). The molecular structure of the 10 top-scored compounds can be seen in **Figure 4**. The three compounds with the highest average binding energy were Dutasteride and Solifenacin, both with − 12.1 kcal/mol energy, followed by Rolapitant with −11.7 kcal/mol energy.

**Figure 4.**
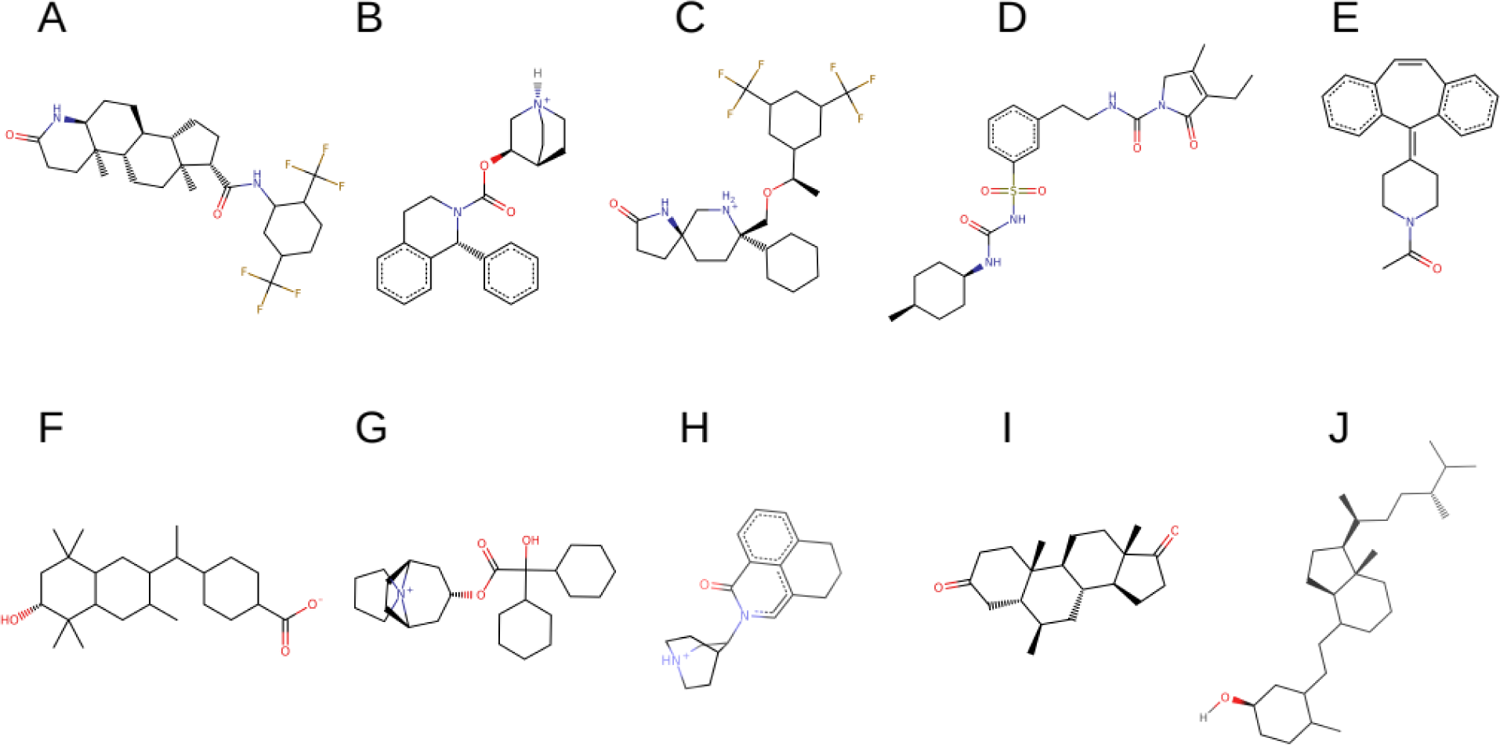
Molecular structure of the 10 top-scored compounds obtained after applying the Structure Based Virtual Screening protocol. The molecular structure is shown as SVG format. The name of each drug and its average binding energy energy (in kcal/mol) is: A) Dutasteride, −12.1; B) Solifenacin, −12.1; C) Rolapitant, − 12.0; D) Glimepiride, −11.9; E) Cyproheptadine, −11.8; F) Bexarotene, −11.7; G) Trospium, −11.4; H) Palonosetron, 11.0, I) Exemestane, −11.0; J) Ergocalciferol, −11.0; C16 (control), −10.4.

**Table 2.** 10 top-scored compounds obtained after applying the whole Structure Based Virtual Screening protocol. Drug name, PubChem ID, drug type and treated condition, and the obtained average Vina energy and Vina docking score energies are shown.

| **Drug name** | **PubChem ID** | **Drug type - treated condition** | **Energy** | **Energy Chain A** | **Energy Chain B** |
| --- | --- | --- | --- | --- | --- |
| **Dutasteride** | 6918296 | 5α-reductase inhibitor – treatment of benign prostatic hyperplasia and | **-12.1** | -11.9 | -12.2 |
| **Solifenacin** | 11155732 (impurity G) | Muscarinic receptor antagonist – treatment of urge urinary incontinence and overactive bladder syndrome | **-12.1** | -11.9 | -12.2 |
| **Rolapitant** | 57758603 (1+ isomer) | NK1 receptor antagonist – prophylaxis of nausea vomiting (antiemetic) | **-12.0** | -12.3 | -11.7 |
| **Glimepiride** | 71317078 (Impurity D) | KATP channel blocker of beta cells – treatment of Diabetes Mellitus Type 2 | **-12.0** | -12.4 | -11.5 |
| **Cyproheptadine** | 524039 (M acetylated) | Serotonin and histamine receptor antagonist – treatment of allergy, | **-11.8** | -12.1 | -11.4 |
| **Bexarotene** | 10067357 (6OH) | RXRs activator – treatment of cutaneous T-cell lymphoma | **-11.7** | -12.1 | -11.2 |
| **Trospium** | 5284632 | Muscarinic receptor antagonist – treatment of urge urinary incontinence and overactive bladder syndrome | **-11.4** | -11.8 | -11.0 |
| **Palonosetron** | 6101833 (3-ene) | Serotonin antagonist blocks 5-HT<sub>3</sub> receptors – prophylaxis of nausea and | **-11.0** | -11.0 | -10.9 |
| **Exemestane** | 60198 | Aromatase inhibitor – treatment of estrogen receptor positive breast cancer | **-11.0** | -11.4 | -10.5 |
| **Ergocalciferol** | 5280793 | Vitamin D<sub>2</sub> – treatment of hypovitaminosis | **-11.0** | -11.2 | -10.7 |
| **C16** | **Control** | **PrfA inhibitor** | **-10.4** | **-11.3** | **-9.4** |

The rationale for doing this additional docking using Autodock Vina, although complemented with docking on chain B of PrfA (site B_I_), accounts for the possibility of potential inaccuracies occurring on PyRx. It must be reminded that in this work Openbabel implemented on PyRx automatically transformed 721 ligands from a 1,345 ligand database into individual PDBQT files. It is known that Openbabel occasionally suffers when handling complex molecules resulting in the generation of PDBQT files with poor physical value. To circumvent this possibility, the 3D SDF files corresponding to the 25 top-scored compounds and their main impurities, metabolites and isomers were extracted from PubChem, processed on Chimera and Autodock Tools, and individually checked as explained in section 2.3 and in [39].

### 3.4 Selection of C16, Dutasteride, Solifenacin and Rolapitant poses for the MD simulations

For the C16 compound, the crystallographic poses that are seen in the 6EXL PDB structure were chosen for the MD simulations. These crystallographic conformations corresponded to pose 1 in chain A (−11.3 kcal/mol) and pose 2 (−9.1 kcal/mol) in chain B. In chain B, pose 1 and pose 2 conformations differed in 0.3 kcal/mol only (**Table 3**). For Dutasteride, pose 1 in chain A (−12.4 kcal/mol) and pose 1 in chain B (−11.8 kcal/mol) were selected. Of note, pose 2 in chain A and pose 2 in chain B were more than 1.0 kcal/mol lower than their corresponding pose 1 conformations (**Table 3**). For Solifenacin, pose 1 in chain B (−12.2 kcal/mol) and pose 2 in chain A (−11.8 kcal/mol) were selected. Whilst pose 1 and pose 2 conformations in chain A differed by only 0.1 kcal/mol, pose 1 and pose 2 conformations in chain B differed by 0.8 kcal/mol (**Table 3**). Therefore, pose 1 in chain B was selected which corresponded to pose 2 in chain A. For Rolapitant, pose 1 in chain B (−11.6 kcal/mol) was selected even though pose 1 in chain A displayed higher energy (−12.3 kcal/mol). However, the corresponding conformations of pose 1 and pose 2 in chain A could not be found in chain B. Consequently, pose 1 in chain B was selected, which corresponded to pose 3 in chain A (−11.0 kcal/mol). Interestingly, pose 1 and pose 2 in chain A displayed the same energy with very minor structural differences (**Table 3**).

**Table 3.**
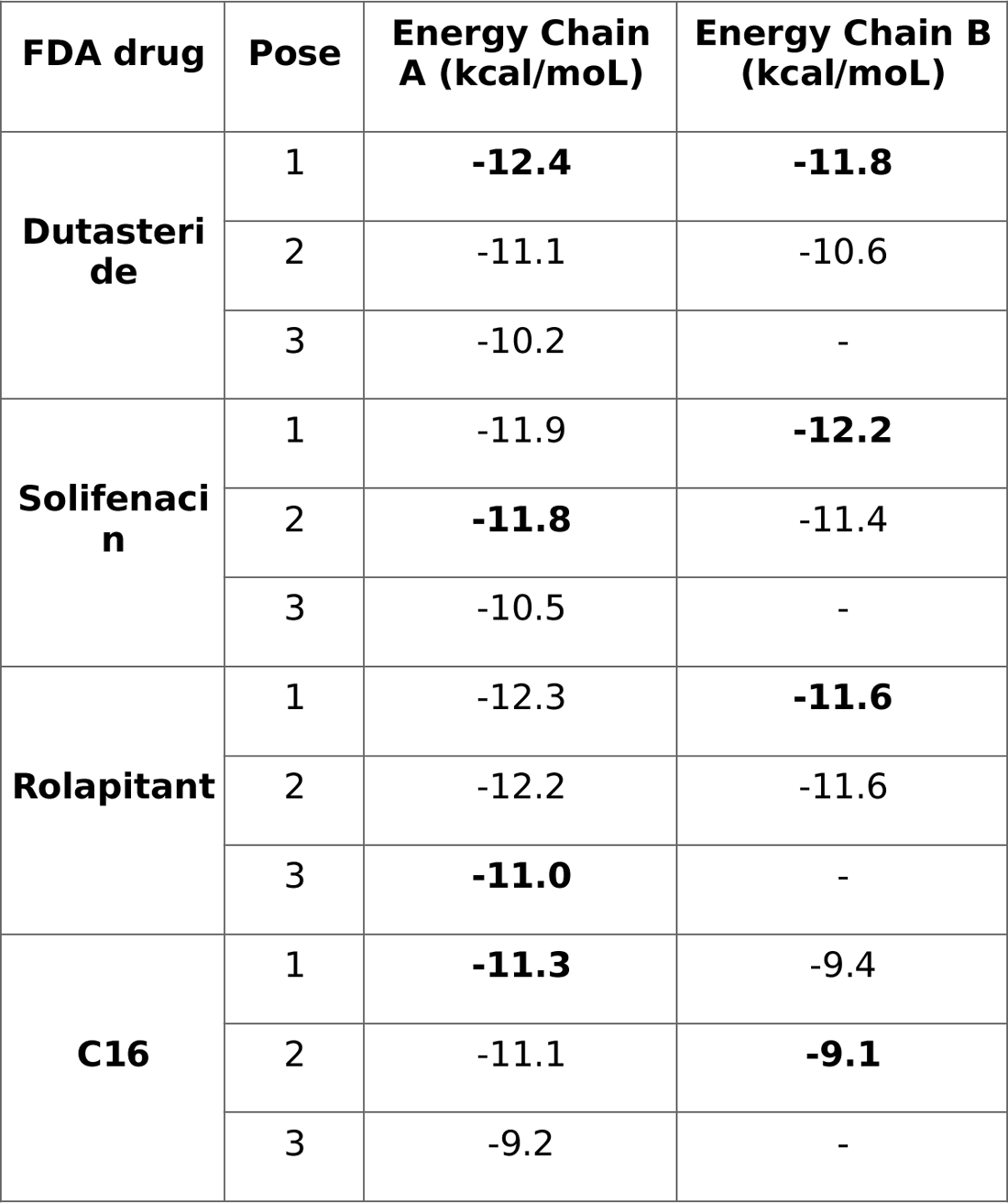
The Vina docking score of the top-3 poses obtained after the final Virtual Screening on Vina for Dutasteride, Solifenacin, Rolapitant and C16 are shown. For the four compounds only two poses were obtained in chain B. The final poses that were chosen for the MD simulation of the PrfA complexes are highlighted in bold. For more information see text.

The three top-scored Vina energy poses for Dutasteride, Solifenacin, Rolapitant and the C16 compound can be seen in **Table 3**. The selected poses can ben seen superimposed in chain A and chain B of PrfA in **Figure 5**.

**Figure 5.**
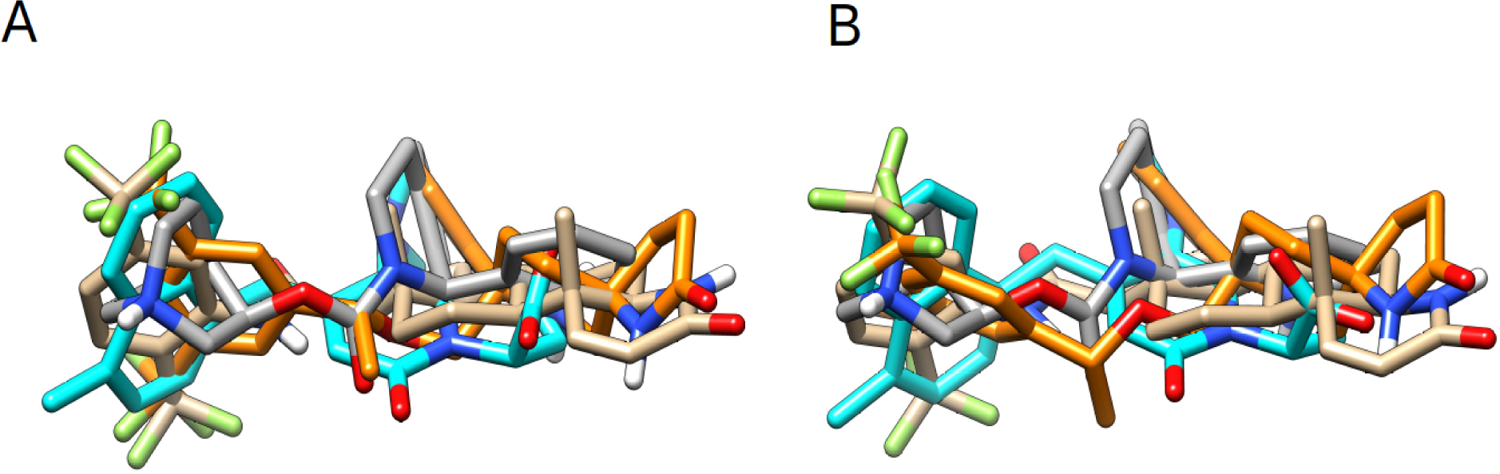
Superimposed poses of C16, Dutasteride, Solifenacin and Rolapitant obtained after performing molecular docking against PrfA’s structure using Autodoc Vina. These poses were chosen for the MD simulations of the compounds complexed to PrfA. Poses in A) Site A_I_ and B) Site B_I_ of PrfA are shown. C16 compound, Dutasteride, Solifenacin and Rolapitant are represented in sticks where C16 is colored in cyan, Dutasteride in brown, Solifenacin in dim grey and Rolapitant in orange. Figures were prepared and rendered on Chimera. For more details see text.

### 3.5 MD simulations of the C16:PrfA complex

In the present study, MD simulations of the C16:PrfA complex were treated as positive controls. Three independent 100-ns MD simulations were performed to check the stability and dynamics of the C16:PrfA complex. The three simulations showed a remarkably high stability for both protein backbone chains (**Figure 6A, 6B, 6C**) and the C16 molecules (**Figure 6D, 6E, 6F**). The average RMSD values obtained for backbone proteins were exactly 1.46 Å, with SD values of 0.15 Å and 0.16 Å for chain A and chain B respectively (**Table 4**). The average RMSD values obtained for the C16 molecules were 0.66 Å and 0.72 Å in chain A and chain B respectively, with SD values of 0.18 Å and 0.16 Å in chain A and chain B respectively (**Table 4**). On the one hand, the low RMSD values obtained for both backbone protein chains (<2.0 Å) and C16 molecules (<1.0 Å) indicate that the starting structures changed very little during the simulated time. On the other hand, the low SD values obtained for both backbone protein chains and C16 molecules (<0.20 Å) indicate that extremely low molecular reorganizations occurred for both backbone protein chains and C16 molecules during the simulations. Overall, the three MD C16:PrfA complex simulations suggest that C16 compound binds PrfA tightly, that the complex is very stable and that very little structural rearrangements are produced once the complex is formed. These results are aligned with those obtained in our previous study [39] and confirm the suitability of treating the C16:PrfA complex as the positive control.

**Figure 6.**
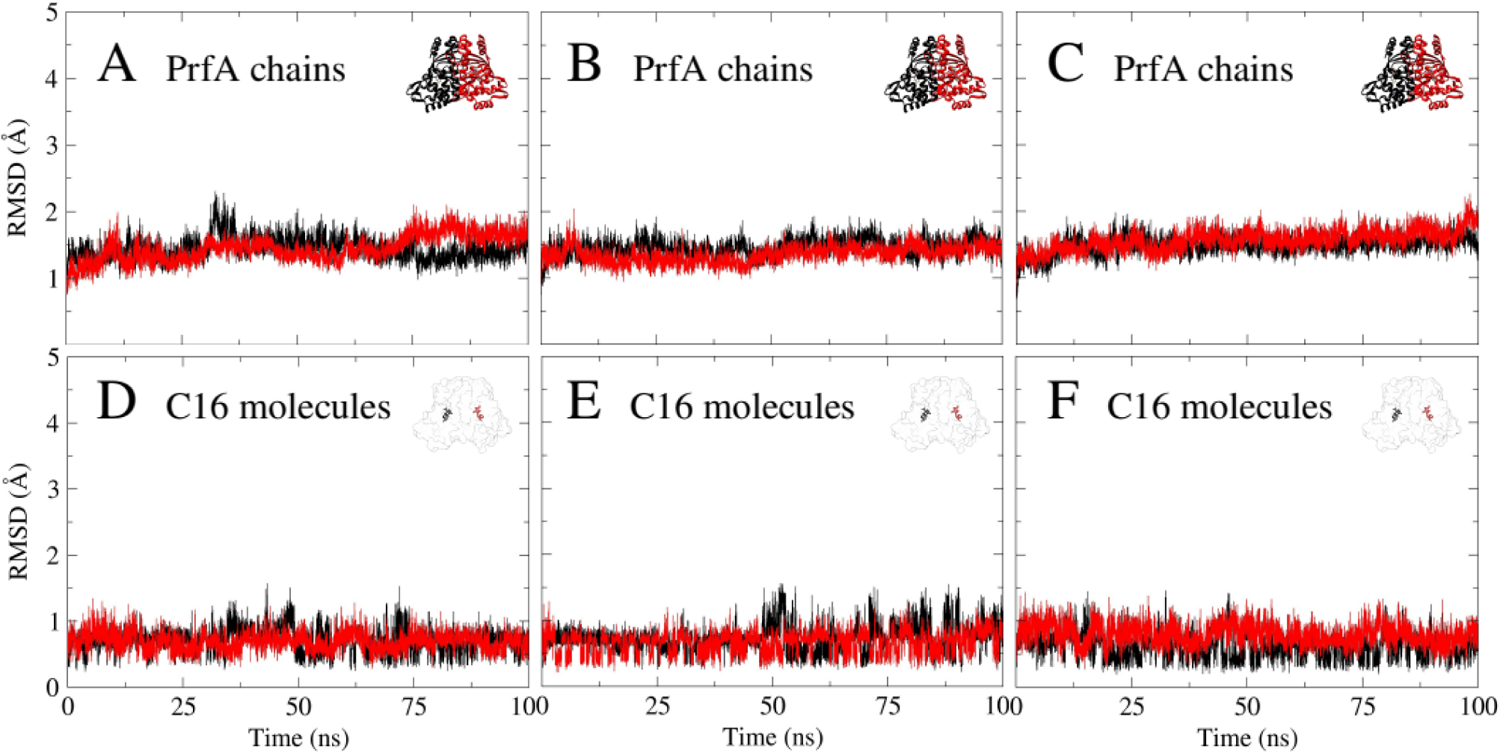
RMSD values obtained for three independent 100-ns MD simulations of PrfA:C16 complex. A-B-C) Backbone RMSD values of PrfA chains and D-E-F) RMSD values of the C16 compounds. A-D) MD simulation 1, B-E) MD simulation 2 and C-F) MD simulation 3. Chain A of PrfA is colored in black and chain B in red. C16 molecule in chain A is colored in black and in chain B is colored in red.

**Table 4.** Average RMSD and SD values calculated for each of the three 100-ns MD simulations of the C16:PrfA complex. Average RMSD and SD values were calculated for the backbone protein chains and the C16 molecules. An average of the calculated average RMSD and SD values is also provided. Value units are shown in Armstrong.

| **PrfA**<br>**com-**<br>**plex** | **Simu-**<br>**lation**<br>**num-**<br>**ber** | **Backbone protein** |  |  |  | **C16 compound molecule** |  |  |  |
| --- | --- | --- | --- | --- | --- | --- | --- | --- | --- |
|  |  | **chain A** |  | **chain B** |  | **chain A** |  | **chain B** |  |
|  |  | **RMSD** | **SD** | **RMSD** | **SD** | **RMSD** | **SD** | **RMSD** | **SD** |
| **PrfA:C1**<br>**6** | **1** | 1.44 | 0.17 | 1.45 | 0.20 | 0.67 | 0.17 | 0.69 | 0.14 |
|  | **2** | 1.45 | 0.13 | 1.35 | 0.13 | 0.72 | 0.20 | 0.66 | 0.17 |
|  | **3** | 1.49 | 0.14 | 1.57 | 0.16 | 0.60 | 0.17 | 0.80 | 0.16 |
|  | **Aver-**<br>**age** | **1.46** | **0.15** | **1.46** | **0.16** | **0.66** | **0.18** | **0.72** | **0.16** |

### 3.6 MD simulations of Dutasteride:PrfA, Solifenacin:PrfA and Rolapitant:PrfA complexes

#### Dutasteride:PrfA

The three 100-ns Dutasteride:PrfA MD simulations showed remarkable high stability of the protein backbone chains (**Figure 7A, 7B, 7C**). The average RMSD values of backbone proteins were 1.67 Å and 1.55 Å for chain A and chain B respectively, with average SD values of 0.17 Å and 0.20 Å for chain A and chain B respectively (**Table 5**). These values are in the same range as those obtained for the C16:PrfA complex (**Table 4**) and thus highlight the high stability of PrfA’s protein chains in the presence of Dutasteride molecules. Overall, the three 100-ns Dutasteride:PrfA MD simulations showed that Dutasteride molecules are stable inside the protein chains (**Figure 7D, 7E, 7F)**. The average RMSD values of Dutasteride molecules were 1.82 Å and 2.27 Å in chain A and chain B respectively, with average SD values of 0.39 Å and 0.26 Å in chain A and chain B respectively (**Table 5**). Whilst stabilization of the Dutasteride molecule in chain A was very fast in simulation 2 (**Figure 7E**), stabilization was slow in simulations 1 (**Figure 7D**) and 3 (**Figure 7F**) and occurred only after the first 12 ns and 24 ns respectively. Once Dutasteride molecules in chain A were stabilized, little fluctuation was seen in simulations 1 (**Figure 7D**) and 2 (**Figure 7E**) but some fluctuation was seen in simulation 3 (**Figure 7F**). Dutasteride molecules in chain B showed little fluctuation in the three simulations (**Figure 7D, 7E, 7F**). Interestingly, Dutasteride molecules in chain B stabilized at RMSD values around 2.5 Å in simulations 1 (**Figure 7D**) and 3 (**Figure 7E**), whilst in simulation 3 (**Figure 7F**), Dutasteride stabilized at around 1.8 Å. Overall, the obtained data shows that PrfA’s protein chains in the presence of Dutasteride molecules are very stable with extremely little fluctuation, at levels comparable to those seen in the C16:PrfA complex. The simulations also show that Dutasteride molecules once stabilized inside PrfA, are quite stable showing very little fluctuation except perhaps for Dutasteride in chain A, where some fluctuation is noted in simulation 3 (**Figure 7F**). Therefore, the data derived from the simulations suggest that PrfA binds Dutasteride and that the complex remains very stable during the simulated time.

**Figure 7.**
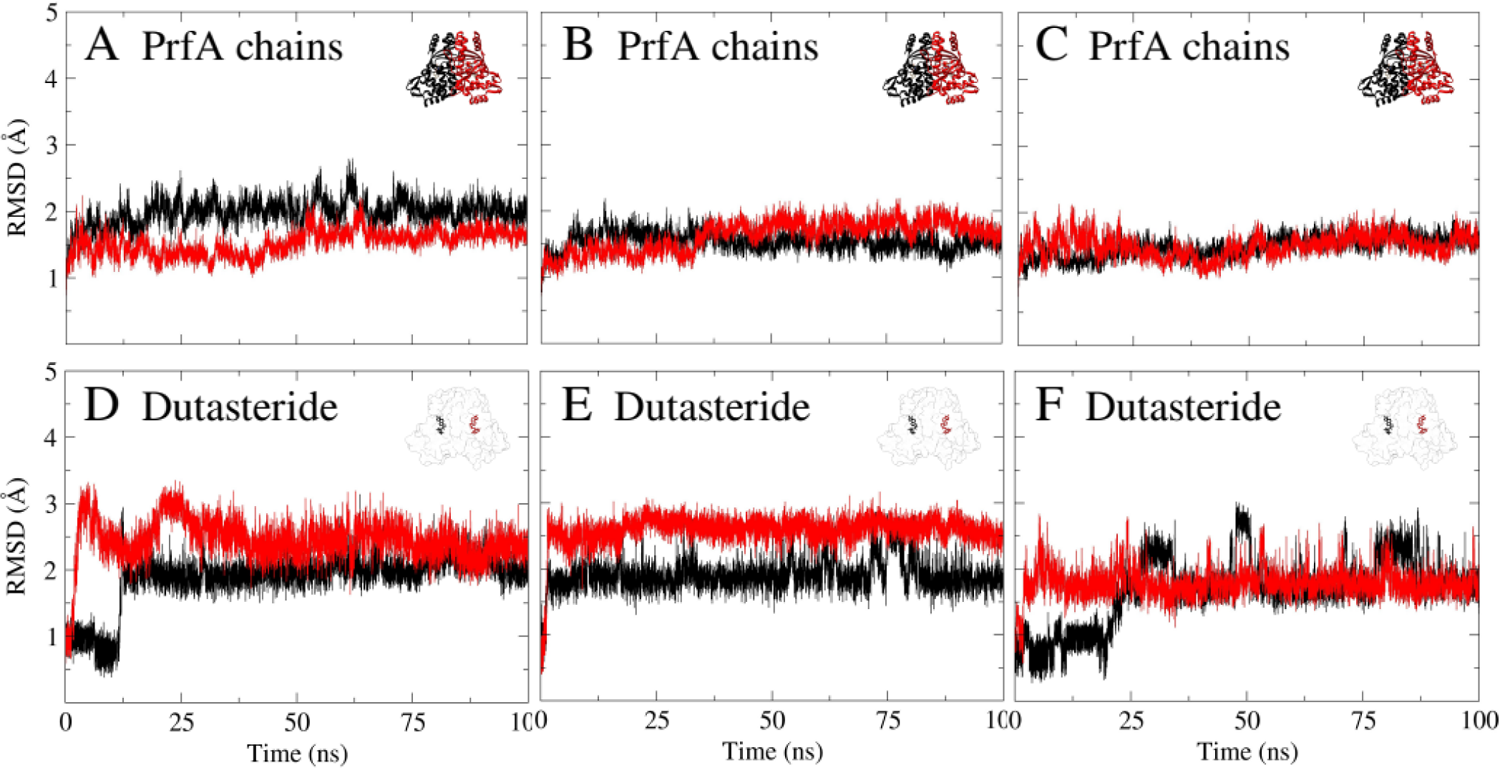
RMSD values obtained for three independent 100-ns MD simulations of PrfA:Dutasteride complex. A-B-C) Backbone RMSD values of PrfA chains and D-E-F) RMSD values of Dutasteride molecules. A-D) MD simulation 1, B-E) MD simulation 2 and C-F) MD simulation 3. Chain A of PrfA is colored in black and chain B in red. Dutasteride molecule in chain A is colored in black and in chain B is colored in red.

**Table 5.**
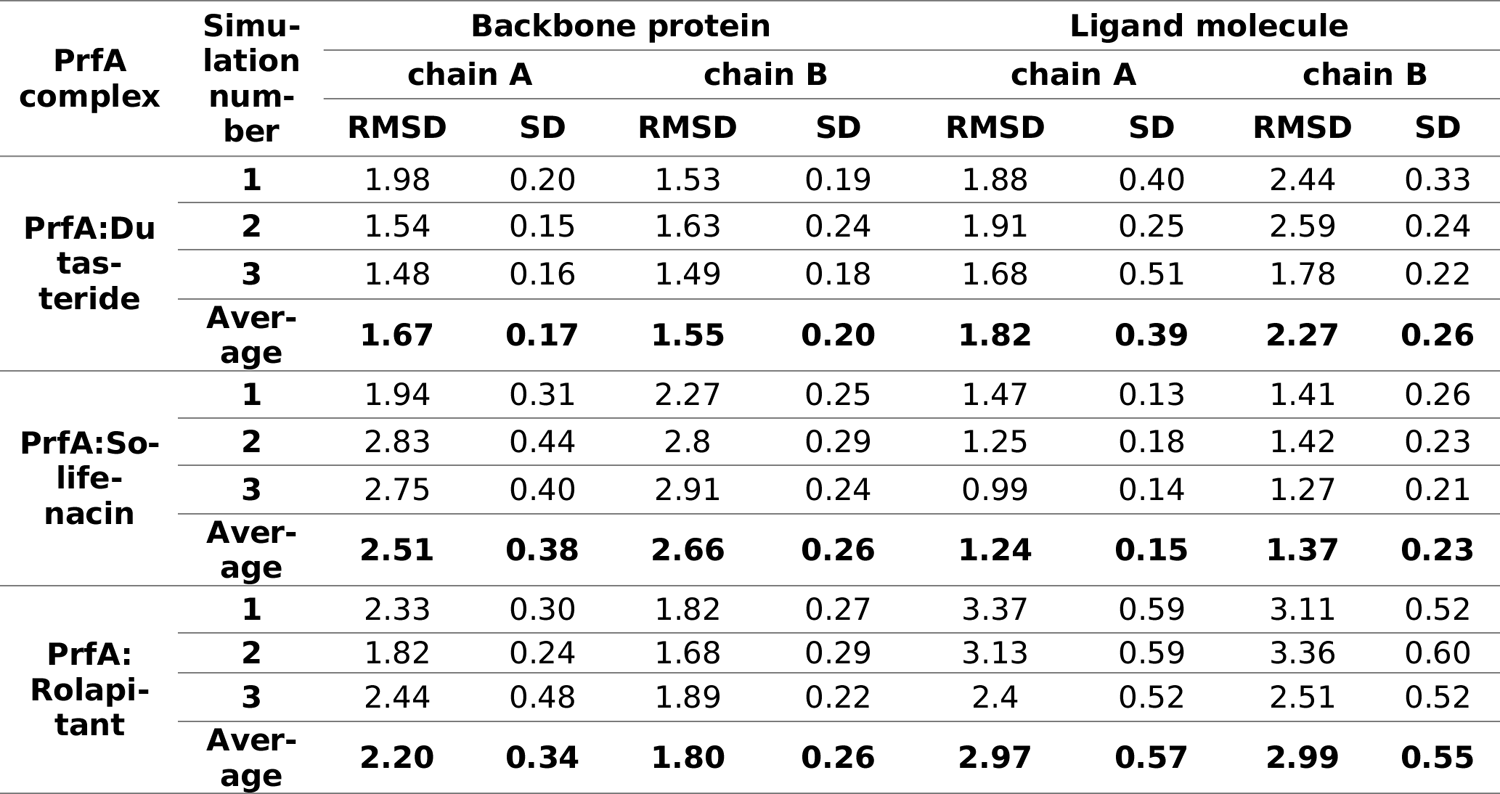
Average RMSD and SD values calculated for each of the three 100-ns MD simulations of Dutasteride:PrfA, Solifenacin:PrfA and Rolapitant:PrfA complexes. Average RMSD and SD values were calculated for the backbone protein chains and the drug molecules. An average of the calculated average RMSD and SD values is also provided. Value units are shown in Armstrong.

#### Solifenacin:PrfA

The three 100-ns Solifenacin:PrfA MD simulations showed moderate stability of the protein backbone chains (**Figure 8A, 8B, 8C**). The average RMSD values of backbone protein chains were 2.51 Å and 2.66 Å for chain A and chain B respectively, with average SD values of 0.38 Å and 0.26 Å for chain A and chain B respectively (**Table 5**). Chain A of PrfA stabilized at RMSD values around 2.8 Å in simulations 2 (**Figure 8B**) and 3 (**Figure 8C**), whilst in simulation 1 (**Figure 8A**) it stabilized at values around 2.2 Å. Chain B showed a more stable behavior and stabilized at RMSD values around 2.8 Å in simulations 2 (**Figure 8B**) and 3 (**Figure 8C**), whilst in simulation 1 (**Figure 8A**) it stabilized at values around 1.9 Å. The three 100-ns Solifenacin:PrfA MD simulations showed very high stability for the Solifenacin molecules both in chain A and chain B (**Figure 8D, 8E, 8F**). The average RMSD values of Solifenacin molecules were 1.24 Å and 1.37 Å in chain A and chain B respectively, with average SD values of 0.15 Å and 0.23 Å in chain A and chain B respectively (**Table 5**). Considering that the number of rotatable bonds of Solifenacin is six while both C16 compound and Dutasteride have four, the obtained low SD values for the Dutasteride molecules, especially in chain A, is remarkable. Even though both protein chains seem to fluctuate slightly, the data suggests that PrfA binds Solifenacin strongly and that the complex is stable. It may be the case that the full equilibrium of the Solifenacin:PrfA complexes may have not reached after 100-ns and therefore, longer simulation times may have been needed to reach full equilibrium.

**Figure 8.**
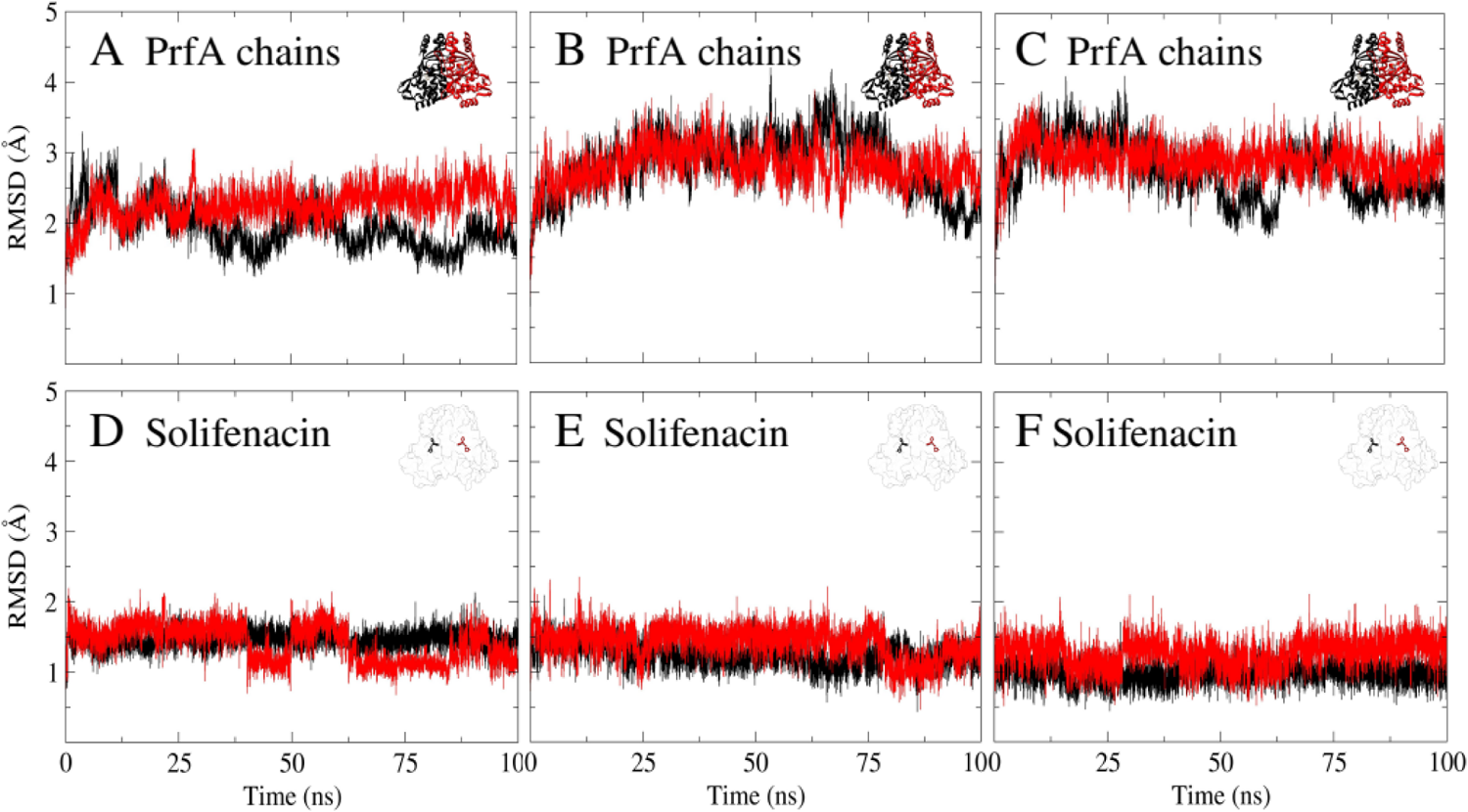
RMSD values obtained for three independent 100-ns MD simulations of PrfA:Solifenacin complex. A-B-C) Backbone RMSD values of PrfA chains and D-E-F) RMSD values of Solifenacin molecules. A-D) MD simulation 1, B-E) MD simulation 2 and C-F) MD simulation 3. Chain A of PrfA is colored in black and chain B in red. Solifenacin molecule in chain A is colored in black and in chain B is colored in red.

#### Rolapitant:PrfA

The three 100-ns Rolapitant:PrfA MD simulations showed poor stability of the protein backbone chains (**Figure 9A, 9B, 9C**). The average RMSD values for backbone proteins were 2.20 Å and 1.80 Å for chain A and chain B respectively, with SD values of 0.34 Å and 0.26 Å for chain A and chain B respectively (**Table 5**). Even though these values are similar to those obtained for the Solifenacin:PrfA complex, a clear positive drift is observed for the protein chains in all simulations (**Figure 9A, 9B, 9C**). The more extreme drift was observed for chain B in simulation 3 (**Figure 9C**), where the starting RMSD values around 1.5 Å ended up above 3.0 Å at the end of the simulation. It is obvious that the Rolapitant:PrfA complex did not reach stabilization during the simulated 100-ns time. The three 100-ns Rolapitant:PrfA MD simulations showed very unstable Rolapitant molecules inside the protein chains (**Figure 9D, 9E, 9F**). The obtained average RMSD values for Rolapitant molecules were 2.97 Å and 2.99 Å in chain A and chain B respectively, with average SD values of 0.57 Å and 0.55 Å in chain A and chain B respectively. The obtained RMSD and SD values for Rolapitant molecules are by far the highest among the four simulated compound complexes (**Table 5**). Overall, the obtained data strongly suggests that Rolapitant interacts weakly with PrfA, if at all. Rolapitant molecules inside PrfA chains appear to be in continuous structural reorganization, which may well be the reason of why the equilibrium is not acquired during the 100-ns MD simulations.

**Figure 9.**
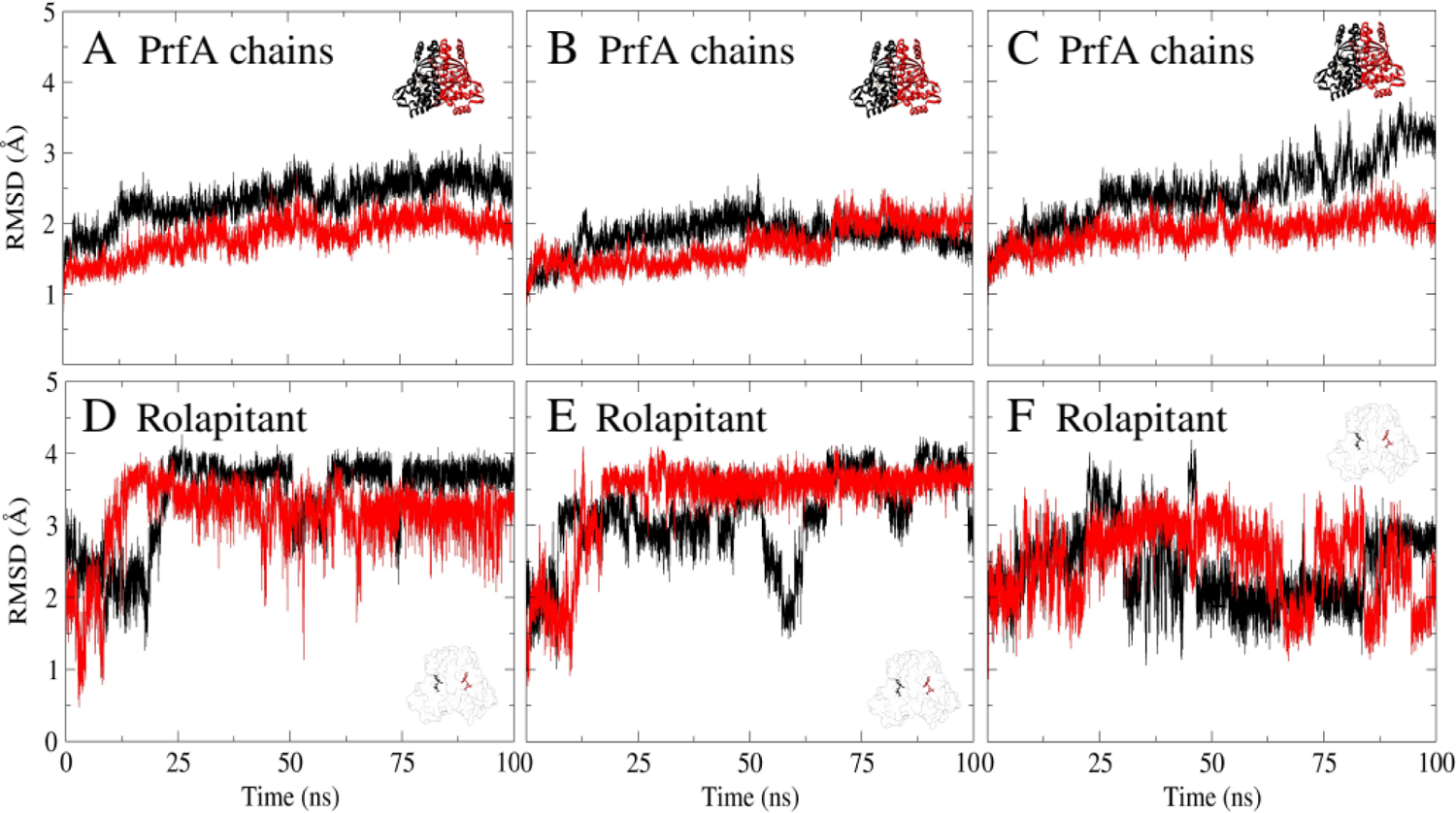
RMSD values obtained for three independent 100-ns MD simulations of PrfA:Rolapitant complex. A-B-C) Backbone RMSD values of PrfA chains and D-E-F) RMSD values of Rolapitant molecules. A-D) MD simulation 1, B-E) MD simulation 2 and C-F) MD simulation 3. Chain A of PrfA is colored in black and chain B in red. Rolapitant molecule in chain A is colored in black and in chain B is colored in red.

## 4. DISCUSSION

The emergence of multi-drug resistant bacteria represents one of the most serious global challenges that healthcare systems will have to face in the coming decades [17]. Although the initial concern was primarily focused on nosocomial bacteria, it has now been extended to all pathogenic bacteria, as multi-drug resistance is becoming prevalent globally. For this reason, the WHO has recently called on all nations to develop novel drugs and treatment methods to fight the rise of this phenomenon [17]. In recent years, antibiotic-resistant *L. monocytogenes* strains have been identified at alarming rates in both clinical and food samples [11]. Several of these antibiotic-resistant *L. monocytogenes* strains have already been confirmed to be multi-drug resistant [12–16]. Considering that the human population is aging and that the elderly are particularly susceptible to listeriosis, a near-future crisis in the management and treatment of invasive listeriosis is anticipated unless novel listeria-specific drugs are developed [5,17]. Current antibiotic therapies against *L. monocytogenes* target its essential biochemical pathways [10]. The gold standard antibiotics in the treatment of listeriosis are penicillins and aminoglycosides [10]. Whilst penicillins target the peptidoglycane cell synthesis, aminoglycosides target the protein synthesis. Contrary, a non-conventional approach that is starting to be considered in the scientific community is to target the virulence of the microorganism rather than the essential and traditional biochemical pathways [35–38]. The underlying idea is to disarm the pathogenicity of the bacteria to such an extent that the pathogens can subsequently be eliminated by the host synergistically with antibiotics or without them. Inhibition of crucial virulence-activating transcriptional factors is already being considered in other pathogenic bacteria [38]. If this strategy were applied to listeriosis, PrfA would be the most promising candidate.

PrfA is a site-specific DNA binding transcriptional regulator whose expression levels are maximized at 37°C, the human body temperature. As a transcriptional factor, PrfA regulates the expression levels of several key virulence factors like Listeriolysin and Acting assembly inducing protein, which are both indispensable for intracellular colonization, among others [26–28]. The first generation of PrfA inhibitors like C01 and C16 were simple ring-fused 2-pyridone compounds. These compounds were shown to decrease virulence factor expression, reduce bacterial uptake into eukaryotic cells, and improve the survival of chicken embryos infected with *L. monocytogenes* [31,32]. Some years later, IWP-2, a Wnt signaling inhibitor, was identified to also inhibit PrfA. IWP-2 was shown to enable lysosomal *L. monocytogenes* clearance from spacious replication vacuoles in infected macrophages [33]. More recently, PS900, a second generation highly substituted ring-fused 2-pyridone compound, has been shown to not only inhibit PrfA but to disrupt *L. monocytogenes*’ general efflux [34].

In the present work, a computational drug repurposing approach has been applied in the hope of identifying FDA drugs with potential PrfA inhibitory activity. One of the main advantages of drug repurposing over traditional drug development is that the compounds being evaluated are approved drugs that are already on the market [20]. Therefore, these molecules are safe to be used in humans [20]. Additionally, the economic and go-to-market readiness of this strategy are of great benefit [20]. For this work, an FDA-approved drugs dataset of small molecule compounds was virtually screened by docking the molecules against the structure of PrfA. Data from the molecular docking and MD simulations suggest that Dutasteride and Solifenacin drugs may bind PrfA, and as a result, may show inhibitory activity against *L. monocytogenes*. Dutasteride, or N-[2,5-bis(trifluoromethyl)phenyl]-3-oxo-4-aza-5α-androst-1-ene-17β-carboxamide, is a synthetic androstane steroid and a 4-azasteroid. Dutasteride is a 5α-reductase inhibitor and therefore, a dihydrotestosterone blocker. Dutasteride is considered an anti-androgenic medication and is principally used in the treatment of enlarged prostate and androgenic alopecia [52,53]. The three 100-ns MD simulations of Dutasteride:PrfA complex show that the PrfA chains are extremely stable in the presence of Dutasteride molecules (**Figure 7A, 7B, 7C**), which may indicate that PrfA binds Dutasteride tightly. Solifenacin, or [(3R)-1-azabicyclo[2.2.2]octan-3-yl] (1S)-1-phenyl-3,4-dihydro-1H-isoquinoline-2-carboxylates, is a muscarinic antagonist with anti-spasmodic properties used to treat urinary incontinence and urinary frequency associated with an overactive bladder [54]. Solifenacin is considered a competitive cholinergic receptor antagonist, selective for the M3 receptor subtype. The three 100-ns MD simulations of Solifenacin:PrA complex show that Solifenacin inside PrfA chains molecules are very stable (**Figure 8D, 8E, 8F**), which suggest that PrfA may also bind Solifenacin strongly.

In conclusion, the rise of multi-drug resistant *L. monocytogenes* strains [11–16] combined with an irreversible aging of the population, will negatively impact invasive listeriosis treatments in the coming decades [17]. Whilst current legislation and surveillance programs have helped reduce the incidence of listeriosis cases in developed countries, modern consumer habits, such as the consumption of RTE foods, are already challenging this progress. In the most recent and deadliest listeriosis outbreak, where more than 200 people died, contaminated RTE processed meat products were confirmed to have caused the outbreak [8]. Therefore, the identification of compounds targeting non-conventional pathways of *L. monocytogenes* will soon become indispensable [17,35,38]. Once identified, these compounds could be developed into fully novel listeria-specific drugs that may be used to treat multi-drug resistant *L. monocytogenes* infections. These novel drugs will help alleviate the foreseeable healthcare crisis expected from the raise of antibiotic resistance [17]. In the present work, several FDA-approved drugs that may bind PrfA have been identified. The molecular docking and MD simulations data suggest that the top two compounds, Dutasteride a Solifenacin, may bind PrfA strongly, and as a result, might inhibit *L. monocytogenes.* The use of Dutasteride and Solifenacin is safe in humans as they are currently prescribed in general medical practice [52–54]. The scientific community is encouraged to experimentally test Dutasteride and Solifenacin drugs against *L. monocytogenes.* If the inhibitory activity is confirmed, their fresh chemical scaffolds will represent valid starting points for the rapid development of novel disruptive listeria-specific drugs.

## 5. CONCLUSIONS

In the present manuscript, several FDA drugs with potential PrfA inhibitory activity are described. The drugs were identified after applying a combined structural-based virtual screening and Molecular Dynamics strategy. According to the obtained data, Dutasteride and Solifenacin are the most promising drug candidates. These drugs might be developed into fully listeria-specific drugs that could one day be used to combat multi-resistant *Listeria monocytogenes* strains.

## Notes

### Competing Interest Statement

The authors have declared no competing interest.

## References

[1] J.A. Vázquez-Boland, M. Kuhn, P. Berche, T. Chakraborty, G. Domínguez-Bernal, W. Goebel, B. González-Zorn, J. Wehland, J. Kreft, Listeria Pathogenesis and Molecular Virulence Determinants, Clin Microbiol Rev. 14 (2001) 584–640. 10.1128/CMR.14.3.584-640.2001.

[2] O. Nwaiwu, What are the recognized species of the genus Listeria?, Access Microbiol. 2 (2020) acmi000153. 10.1099/acmi.0.000153.

[3] K.-H. Byun, H.J. Kim, Survival strategies of Listeria monocytogenes to environmental hostile stress: biofilm formation and stress responses, Food Sci Biotechnol. 32 (2023) 1631–1651. 10.1007/s10068-023-01427-6.

[4] O. Disson, M. Lecuit, Targeting of the central nervous system by Listeria monocytogenes, Virulence. 3 (2012) 213–221. 10.4161/viru.19586.

[5] P. Muñoz, L. Rojas, E. Bunsow, E. Saez, L. Sánchez-Cambronero, L. Alcalá, M. Rodríguez-Creixems, E. Bouza, Listeriosis: An emerging public health problem especially among the elderly, Journal of Infection. 64 (2012) 19–33. 10.1016/j.jinf.2011.10.006.

6. FDA, Listeria (Listeriosis) - https://www.fda.gov/food/foodborne-pathogens/listeria-listeriosis, FDA. (2023). https://www.fda.gov/food/foodborne-pathogens/listeria-listeriosis (accessed October 29, 2023).

[7] E.F.S. Authority, European Centre for Disease Prevention and Control, The European Union One Health 2021 Zoonoses Report, EFSA Journal. 20 (2022) e07666. 10.2903/j.efsa.2022.7666.

[8] A.M. Smith, N.P. Tau, S.L. Smouse, M. Allam, A. Ismail, N.R. Ramalwa, B. Disenyeng, M. Ngomane, J. Thomas, Outbreak of Listeria monocytogenes in South Africa, 2017-2018: Laboratory Activities and Experiences Associated with Whole-Genome Sequencing Analysis of Isolates, Foodborne Pathog Dis. 16 (2019) 524–530. 10.1089/fpd.2018.2586.

[9] V. Goulet, L.A. King, V. Vaillant, H. de Valk, What is the incubation period for listeriosis?, BMC Infect Dis. 13 (2013) 11. 10.1186/1471-2334-13-11.

[10] S. Thønnings, J.D. Knudsen, H.C. Schønheyder, M. Søgaard, M. Arpi, K.O. Gradel, C. Østergaard, Danish Collaborative Bacteraemia Network (DACOBAN), Antibiotic treatment and mortality in patients with Listeria monocytogenes meningitis or bacteraemia, Clin Microbiol Infect. 22 (2016) 725–730. 10.1016/j.cmi.2016.06.006.

[11] A.N. Olaimat, M.A. Al-Holy, H.M. Shahbaz, A.A. Al-Nabulsi, M.H. Abu Ghoush, T.M. Osaili, M.M. Ayyash, R.A. Holley, Emergence of Antibiotic Resistance in Listeria monocytogenes Isolated from Food Products: A Comprehensive Review, Compr Rev Food Sci Food Saf. 17 (2018) 1277–1292. 10.1111/1541-4337.12387.

[12] L. Mpondo, K.E. Ebomah, A.I. Okoh, Multidrug-Resistant Listeria Species Shows Abundance in Environmental Waters of a Key District Municipality in South Africa, Int J Environ Res Public Health. 18 (2021) 481. 10.3390/ijerph18020481.

[13] A.A. Fallah, S.S. Saei-Dehkordi, M. Mahzounieh, Occurrence and antibiotic resistance profiles of Listeria monocytogenes isolated from seafood products and market and processing environments in Iran, Food Control. 34 (2013) 630–636. 10.1016/j.foodcont.2013.06.015.

[14] E. Abdollahzadeh, S.M. Ojagh, H. Hosseini, E.A. Ghaemi, G. Irajian, M. Naghizadeh Heidarlo, Antimicrobial resistance of Listeria monocytogenes isolated from seafood and humans in Iran, Microbial Pathogenesis. 100 (2016) 70–74. 10.1016/j.micpath.2016.09.012.

[15] Y. Zhang, E. Yeh, G. Hall, J. Cripe, A.A. Bhagwat, J. Meng, Characterization of Listeria monocytogenes isolated from retail foods, International Journal of Food Microbiology. 113 (2007) 47–53. 10.1016/j.ijfoodmicro.2006.07.010.

[16] I. Sakaridis, N. Soultos, E. Iossifidou, A. Papa, I. Ambrosiadis, P. Koidis, Prevalence and Antimicrobial Resistance of Listeria monocytogenes Isolated in Chicken Slaughterhouses in Northern Greece, Journal of Food Protection. 74 (2011) 1017–1021. 10.4315/0362-028X.JFP-10-545.

17. WHO, Global action plan on antimicrobial resistance, (2015). https://www.who.int/publications-detail-redirect/9789241509763 (accessed October 25, 2023).

[18] J. Hughes, S. Rees, S. Kalindjian, K. Philpott, Principles of early drug discovery, Br J Pharmacol. 162 (2011) 1239–1249. 10.1111/j.1476-5381.2010.01127.x.

[19] S. Kraljevic, P.J. Stambrook, K. Pavelic, Accelerating drug discovery, EMBO Rep. 5 (2004) 837–842. 10.1038/sj.embor.7400236.

[20] N. Rao, T. Poojari, C. Poojary, R. Sande, S. Sawant, Drug Repurposing: a Shortcut to New Biological Entities, Pharm Chem J. 56 (2022) 1203–1214. 10.1007/s11094-022-02778-w.

[21] L. Vangeel, W. Chiu, S. De Jonghe, P. Maes, B. Slechten, J. Raymenants, E. André, P. Leyssen, J. Neyts, D. Jochmans, Remdesivir, Molnupiravir and Nirmatrelvir remain active against SARS-CoV-2 Omicron and other variants of concern, Antiviral Research. 198 (2022) 105252. 10.1016/j.antiviral.2022.105252.

22. WHO, Therapeutics and COVID-19: living guideline, (2023). https://www.who.int/publications-detail-redirect/WHO-2019-nCoV-therapeutics-2022.4 (accessed October 29, 2023).

[23] D.R. Owen, C.M.N. Allerton, A.S. Anderson, L. Aschenbrenner, M. Avery, S. Berritt, B. Boras, R.D. Cardin, A. Carlo, K.J. Coffman, A. Dantonio, L. Di, H. Eng, R. Ferre, K.S. Gajiwala, S.A. Gibson, S.E. Greasley, B.L. Hurst, E.P. Kadar, A.S. Kalgutkar, J.C. Lee, J. Lee, W. Liu, S.W. Mason, S. Noell, J.J. Novak, R.S. Obach, K. Ogilvie, N.C. Patel, M. Pettersson, D.K. Rai, M.R. Reese, M.F. Sammons, J.G. Sathish, R.S.P. Singh, C.M. Steppan, A.E. Stewart, J.B. Tuttle, L. Updyke, P.R. Verhoest, L. Wei, Q. Yang, Y. Zhu, An oral SARS-CoV-2 Mpro inhibitor clinical candidate for the treatment of COVID-19, Science. 374 (2021) 1586–1593. 10.1126/science.abl4784.

[24] A.J. Pruijssers, A.S. George, A. Schäfer, S.R. Leist, L.E. Gralinksi, K.H. Dinnon, B.L. Yount, M.L. Agostini, L.J. Stevens, J.D. Chappell, X. Lu, T.M. Hughes, K. Gully, D.R. Martinez, A.J. Brown, R.L. Graham, J.K. Perry, V. Du Pont, J. Pitts, B. Ma, D. Babusis, E. Murakami, J.Y. Feng, J.P. Bilello, D.P. Porter, T. Cihlar, R.S. Baric, M.R. Denison, T.P. Sheahan, Remdesivir Inhibits SARS-CoV-2 in Human Lung Cells and Chimeric SARS-CoV Expressing the SARS-CoV-2 RNA Polymerase in Mice, Cell Rep. 32 (2020) 107940. 10.1016/j.celrep.2020.107940.

[25] F. Kabinger, C. Stiller, J. Schmitzová, C. Dienemann, G. Kokic, H.S. Hillen, C. Höbartner, P. Cramer, Mechanism of molnupiravir-induced SARS-CoV-2 mutagenesis, Nat Struct Mol Biol. 28 (2021) 740–746. 10.1038/s41594-021-00651-0.

[26] A. de las Heras, R.J. Cain, M.K. Bielecka, J.A. Vázquez-Boland, Regulation of Listeria virulence: PrfA master and commander, Current Opinion in Microbiology. 14 (2011) 118–127. 10.1016/j.mib.2011.01.005.

[27] N.E. Freitag, L. Rong, D.A. Portnoy, Regulation of the prfA transcriptional activator of Listeria monocytogenes: multiple promoter elements contribute to intracellular growth and cell-to-cell spread., Infect Immun. 61 (1993) 2537–2544.

[28] T. Chakraborty, M. Leimeister-Wächter, E. Domann, M. Hartl, W. Goebel, T. Nichterlein, S. Notermans, Coordinate regulation of virulence genes in Listeria monocytogenes requires the product of the prfA gene., J Bacteriol. 174 (1992) 568–574.

[29] M. Hall, C. Grundström, A. Begum, M.J. Lindberg, U.H. Sauer, F. Almqvist, J. Johansson, A.E. Sauer-Eriksson, Structural basis for glutathione-mediated activation of the virulence regulatory protein PrfA in Listeria, PNAS. 113 (2016) 14733–14738. 10.1073/pnas.1614028114.

[30] M.L. Reniere, A.T. Whiteley, K.L. Hamilton, S.M. John, P. Lauer, R.G. Brennan, D.A. Portnoy, Glutathione activates virulence gene expression of an intracellular pathogen, Nature. 517 (2015) 170–173. 10.1038/nature14029.

[31] J.A.D. Good, C. Andersson, S. Hansen, J. Wall, K.S. Krishnan, A. Begum, C. Grundström, M.S. Niemiec, K. Vaitkevicius, E. Chorell, P. Wittung-Stafshede, U.H. Sauer, A.E. Sauer-Eriksson, F. Almqvist, J. Johansson, Attenuating Listeria monocytogenes Virulence by Targeting the Regulatory Protein PrfA, Cell Chem Biol. 23 (2016) 404–414. 10.1016/j.chembiol.2016.02.013.

[32] M. Kulén, M. Lindgren, S. Hansen, A.G. Cairns, C. Grundström, A. Begum, I. van der Lingen, K. Brännström, M. Hall, U.H. Sauer, J. Johansson, A.E. Sauer-Eriksson, F. Almqvist, Structure-Based Design of Inhibitors Targeting PrfA, the Master Virulence Regulator of Listeria monocytogenes, J. Med. Chem. 61 (2018) 4165–4175. 10.1021/acs.jmedchem.8b00289.

[33] T.T. Tran, C.D. Mathmann, M. Gatica-Andrades, R.F. Rollo, M. Oelker, J.K. Ljungberg, T.T.K. Nguyen, A. Zamoshnikova, L.K. Kummari, O.J.K. Wyer, K.M. Irvine, J. Melo-Bolívar, A. Gross, D. Brown, J.Y.W. Mak, D.P. Fairlie, K.A. Hansford, M.A. Cooper, R. Giri, V. Schreiber, S.R. Joseph, F. Simpson, T.C. Barnett, J. Johansson, W. Dankers, J. Harris, T.J. Wells, R. Kapetanovic, M.J. Sweet, E.A. Latomanski, H.J. Newton, R.J.R. Guérillot, A. Hachani, T.P. Stinear, S.Y. Ong, Y. Chandran, E.L. Hartland, B. Kobe, J.L. Stow, A.E. Sauer-Eriksson, J. Begun, J.C. Kling, A. Blumenthal, Inhibition of the master regulator of Listeria monocytogenes virulence enables bacterial clearance from spacious replication vacuoles in infected macrophages, PLoS Pathog. 18 (2022) e1010166. 10.1371/journal.ppat.1010166.

[34] R. Dehbanipour, Z. Ghalavand, Anti-virulence therapeutic strategies against bacterial infections: recent advances, Germs. 12 (2022) 262–275. 10.18683/germs.2022.1328.

[35] A.E. Clatworthy, E. Pierson, D.T. Hung, Targeting virulence: a new paradigm for antimicrobial therapy, Nat Chem Biol. 3 (2007) 541–548. 10.1038/nchembio.2007.24.

[36] D.A. Rasko, V. Sperandio, Anti-virulence strategies to combat bacteria-mediated disease, Nat Rev Drug Discov. 9 (2010) 117–128. 10.1038/nrd3013.

[37] S. Mühlen, P. Dersch, Anti-virulence Strategies to Target Bacterial Infections, Curr Top Microbiol Immunol. 398 (2016) 147–183. 10.1007/82_2015_490.

[38] B. Nizami, W. Tan, X. Arias-Moreno, In silico identification of novel PrfA inhibitors to fight listeriosis: A virtual screening and molecular dynamics studies, J Mol Graph Model. 101 (2020) 107728. 10.1016/j.jmgm.2020.107728.

[39] H.M. Berman, J. Westbrook, Z. Feng, G. Gilliland, T.N. Bhat, H. Weissig, I.N. Shindyalov, P.E. Bourne, The Protein Data Bank, Nucleic Acids Research. 28 (2000) 235–242. 10.1093/nar/28.1.235.

[40] G.M. Morris, R. Huey, W. Lindstrom, M.F. Sanner, R.K. Belew, D.S. Goodsell, A.J. Olson, AutoDock4 and AutoDockTools4: Automated Docking with Selective Receptor Flexibility, J Comput Chem. 30 (2009) 2785–2791. 10.1002/jcc.21256.

[41] S. Dallakyan, A.J. Olson, Small-molecule library screening by docking with PyRx, Methods Mol Biol. 1263 (2015) 243–250. 10.1007/978-1-4939-2269-7_19.

[42] O. Trott, A.J. Olson, AutoDock Vina: improving the speed and accuracy of docking with a new scoring function, efficient optimization and multithreading, Journal of Computational Chemistry. 31 (2010) 455. 10.1002/jcc.21334.

[43] T. Sterling, J.J. Irwin, ZINC 15 – Ligand Discovery for Everyone, J. Chem. Inf. Model. 55 (2015) 2324–2337. 10.1021/acs.jcim.5b00559.

[44] W. Humphrey, A. Dalke, K. Schulten, VMD: visual molecular dynamics, J Mol Graph. 14 (1996) 33–38, 27–28. 10.1016/0263-7855(96)00018-5.

[45] S. Kim, J. Chen, T. Cheng, A. Gindulyte, J. He, S. He, Q. Li, B.A. Shoemaker, P.A. Thiessen, B. Yu, L. Zaslavsky, J. Zhang, E.E. Bolton, PubChem 2023 update, Nucleic Acids Research. 51 (2023) D1373–D1380. 10.1093/nar/gkac956.

[46] N.M. O’Boyle, M. Banck, C.A. James, C. Morley, T. Vandermeersch, G.R. Hutchison, Open Babel: An open chemical toolbox, Journal of Cheminformatics. 3 (2011) 33. 10.1186/1758-2946-3-33.

[47] E.F. Pettersen, T.D. Goddard, C.C. Huang, G.S. Couch, D.M. Greenblatt, E.C. Meng, T.E. Ferrin, UCSF Chimera--a visualization system for exploratory research and analysis, J Comput Chem. 25 (2004) 1605–1612. 10.1002/jcc.20084.

48. Chemaxon https://www.chemaxon.com – November 2023, (n.d.).

[49] M.J. Abraham, T. Murtola, R. Schulz, S. Páll, J.C. Smith, B. Hess, E. Lindahl, GROMACS: High performance molecular simulations through multi-level parallelism from laptops to supercomputers, SoftwareX. 1–2 (2015) 19–25. 10.1016/j.softx.2015.06.001.

[50] V. Zoete, M.A. Cuendet, A. Grosdidier, O. Michielin, SwissParam: a fast force field generation tool for small organic molecules, J Comput Chem. 32 (2011) 2359–2368. 10.1002/jcc.21816.

[51] H. Tükenmez, P. Singh, S. Sarkar, M. Çakır, A.H. Oliveira, C. Lindgren, K. Vaitkevicius, M. Bonde, A.E. Sauer-Eriksson, F. Almqvist, J. Johansson, A Highly Substituted Ring-Fused 2-Pyridone Compound Targeting PrfA and the Efflux Regulator BrtA in Listeria monocytogenes, mBio. 14 (2023) e0044923. 10.1128/mbio.00449-23.

[52] T. Arif, K. Dorjay, M. Adil, M. Sami, Dutasteride in Androgenetic Alopecia: An Update, Curr Clin Pharmacol. 12 (2017) 31–35. 10.2174/1574884712666170310111125.

[53] C. Wu, A. Kapoor, Dutasteride for the treatment of benign prostatic hyperplasia, Expert Opin Pharmacother. 14 (2013) 1399–1408. 10.1517/14656566.2013.797965.

[54] J.C. Santos, E.R.Z. Telo, Solifenacin: scientific evidence in the treatment of overactive bladder, Arch Esp Urol. 63 (2010) 197–213.

